# Structural Interactions of Ankyrin B with NrCAM and β_2_ Spectrin

**DOI:** 10.1101/2025.04.30.651523

**Authors:** Venkata R. Chirasani, Victoria A. Haberman, Erik N. Oldre, Barrett D. Webb, Ernest B. Pereira, Wonsuk Yang, Patricia F. Maness

## Abstract

*Ank2* is a high confidence autism spectrum disorder (ASD) gene encoding the spectrin-actin scaffold protein Ankyrin B (AnkB). The 220 kDal isoform of AnkB has multiple functions including developmental spine pruning through L1 family cell adhesion molecules (L1-CAMs) and class 3 Semaphorins on dendrites of pyramidal neurons to achieve an appropriate excitatory balance in the neocortex. Molecular modeling employing AlphaFold was used to predict the structure and interactions of AnkB with the cytoplasmic domain of Neuron-glial Related L1-CAM (NrCAM), and with β2-Spectrin. The validity of the models was assessed by analyzing protein-protein interactions by co-immunoprecipitation from HEK293 cell lysates after mutating key residues in AnkB predicted to impair these associations. Results revealed a pocket with critical residues in the AnkB membrane-binding domain that engages NrCAM at the conserved cytoplasmic motif FIGQY. Alphafold modeling of the AnkB/β2-Spectrin complex also identified key interactions between the AnkB spectrin-binding domain and β2-Spectrin repeats 14-15. Selected ASD-linked mutations in AnkB predicted to impact binding to NrCAM or β2-Spectrin were then assayed for protein interactions. Maternally inherited ASD missense mutations AnkB A368G located in the NrCAM binding pocket and AnkB R977Q in the Zu51 subdomain disrupted associations with NrCAM and β2-Spectrin, respectively. Moreover, AnkB A368G impaired the neuronal function of 220 kDal AnkB in Semaphorin 3F-induced spine pruning in mouse cortical neuron cultures. These new findings provide structural insights into the L1-CAM/AnkB complex and the molecular basis of ASD etiology associated with AnkB missense mutations.

## Introduction

Large scale genomic studies have revealed a diversity of genetic mutations that increase disease risk in autism spectrum disorder (ASD) and other neurodevelopmental diseases (1–3). Many ASD risk genes are linked to altered development of pyramidal neurons (4,5), principal neurons whose dendritic spines harbor the vast majority of excitatory synapses in the neocortex. Cortical hyper-expansion and brain enlargement are vulnerabilities for ASD that become apparent in childhood at the end of year 1 (6–8). Abnormal social and cognitive functions characteristic of ASD are not present at this time, but arise after relevant neural circuits are formed and synaptic connections are refined (9). During normal development dendritic spines and excitatory synapses are initially overproduced, eliminated in substantial numbers in juveniles, and stabilized in adulthood (10–13). In ASD and Fragile X Syndrome, elevated spine density is evident in the adult prefrontal cortex (PFC) (14–18), where essential circuits contribute to social and cognitive function (19). Defective pruning of overproduced spines is therefore a cogent hypothesis for increased spine density and behavioral deficits associated with ASD.

We identified a molecular mechanism during postnatal development of the mouse prefrontal cortex, in which class 3 Semaphorins (Sema3s) drive activity-dependent spine elimination on pyramidal neurons (20). Sema3s bind receptor complexes comprising L1-CAMs and their coreceptors Neuropilin-1/2 (Npn-1/2) and PlexinA1-4 (PlexA1-4) (21–24) (Fig.1A). NrCAM (Neuron-glial Related-CAM) is an L1 family member, which together with Npn-2 and PlexA3, forms a holoreceptor for Sema3F, while CHL1 (Close Homolog of L1) with Npn-2 and PlexA4, forms a holoreceptor for Sema3B mediating spine pruning (22,23). This specificity allows Sema3F to prune spine subpopulations through NrCAM, while Sema3B prunes a distinct spine subpopulation through CHL1. L1, the prototype of the L1-CAM family, binds Npn-1, a coreceptor for Sema3A (25), and also regulates spine density (26). Selective spine pruning through L1-CAMs occurs only on apical, not basal, dendrites of cortical pyramidal neurons. Apical and basal dendrites are differentially specialized for propagating electrical signals and synaptic connectivity (see refs in (27)).

**Figure 1:**
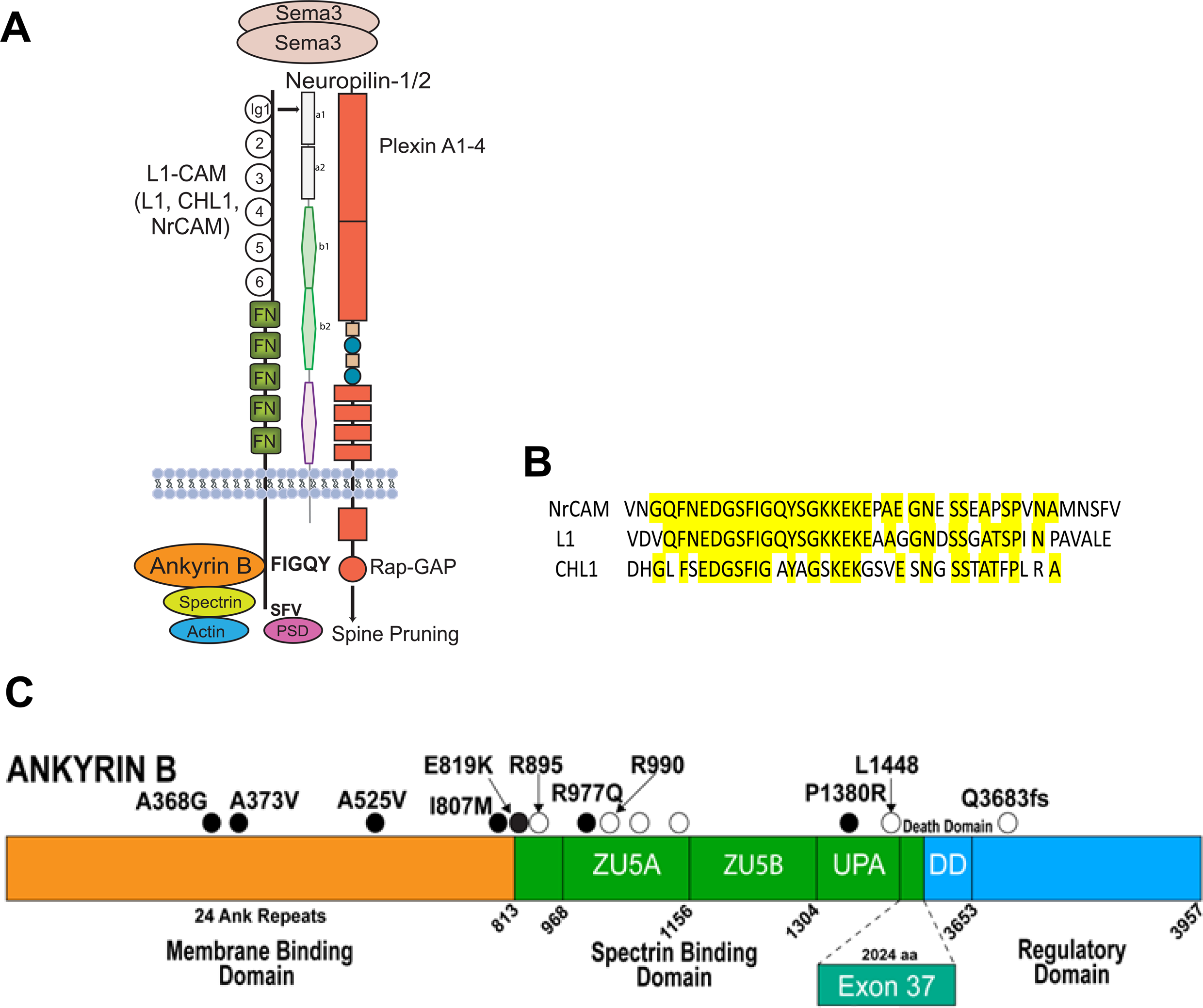
Semaphorin 3 Holoreceptor Complex with Conserved L1-CAM Cytoplasmic Domains, and AnkB Domain Structure with Sites of Human ASD Mutations A. Scheme of secreted class 3 Semaphorins (Sema3 dimers) bound to holoreceptor complex of L1-CAMs (L1, CHL1, NrCAM), Neuropilins (Npn-1/2), and PlexinA1-4. The Ig1 domain of L1-CAMs binds to the a1 domain of Npn1.2. L1-CAM cytoplasmic domains bind to the Spectrin-Actin adaptor AnkB at motif FIGQY, and to PSD proteins at the C-terminal motif SFV. Downstream signaling through the intrinsic Rap-GAP activity of PlexinA culminates in dendritic spine pruning in neurons. B. Sequences of the conserved cytoplasmic domains of NrCAM, L1, and CHL1. Alignment shows identical residues in yellow, including the Ankyrin binding motif FIGQ/AY and flanking residues. C. AnkB-220 domains with sites of human ASD mutations: ● missense; ○ nonsense, frameshift and deletion (fs). The AnkB membrane binding domain (MBD) contains 24 ANK repeats. The spectrin binding domain (SBD) includes ZU5A/B and UPA subdomains. The death domain (DD) and regulatory domain are located in the C-terminal region. A large (2024 amino acid) insert encoded by exon 37 of the *Ank2* gene generates the AnkB-440 splice variant.

All L1-CAMs bind Ankyrin B (AnkB), a spectrin-actin adaptor protein that is encoded by the high confidence ASD risk gene *Ank2* (category 1, SFARI Gene Database) (Fig. 1C). There are two principal AnkB spliced isoforms in the brain (220-and 440-kDa) (28). AnkB-220 is localized to the somatodendritic compartment of neurons (29) and mediates Sema3-induced spine pruning (30), whereas AnkB-440, which contains a large insert encoded by exon 37, is axonal and suppresses axon branching (31,32)(Fig. 1C). L1-CAMs engage Ankyrins through the highly conserved L1-CAM cytoplasmic domain, of which a critical region includes the motif FIGQ/AY (Fig. 1B) (33–35). Mutation of FIGQY to FIGQH in L1 is associated with a human intellectual disability syndrome with varying degrees of hydrocephalus, aphasia, and spastic paraplegia (36,37). L1-FIGQH knock-in mouse models are defective in AnkB binding and exhibit increased spine density on apical dendrites (26), axon guidance errors (38), and unstable interneuron synapses (39,40). Global L1 knockout mice also display increased spine density (26), enlarged brain ventricles, and aberrant axon guidance (41–43).

L1-CAMs bind to the Ankyrin N-terminal membrane-binding domain (MBD), which comprises 24 ANK repeats that form a superhelical solenoid structure (44) (Fig. 1C). The Ankyrin MBD also binds voltage-gated ion channels such as Nav1.2 (29). Binding sites for Nav1.2 and another L1-CAM, Neurofascin (160 kD), are distributed across ANK repeats and partially overlap but are not fully defined (44). Spectrins are Actin-binding proteins comprising elongated heterotetramers of 2 α and 2 β subunits (45). Ankyrins engage β-Spectrins at the spectrin-binding domain (SBD) supermodule Zu5A/B, UPA (Fig.1C), linking Ankyrin-receptor complexes to the actin cytoskeleton (46). The C-terminal region of AnkB contains a death domain and regulatory sequence (Fig. 1C). Human *de novo* and inherited ASD variants have been reported across the *Ank2* sequence of both principal isoforms of AnkB (Fig. 1C) (2,47–52). These variants include missense and truncating mutations in the AnkB MBD and SBD common to both isoforms. Some genetic variants in β2-Spectrin are also associated with autism-like neurodevelopmental syndromes (45,53,54). The molecular interactions between the L1-CAM cytoplasmic domain and AnkB MBD, and between β-Spectrin and the AnkB SBD have not been defined in detail, in part due to the challenge of analyzing their extended protein architectures.

Here we used a molecular modeling and biochemical approach to predict and validate the structural interactions of AnkB with the conserved cytoplasmic domain of NrCAM, and with β2-Spectrin. We generated selected mutations in the AnkB MBD and SBD, and assessed their impact on binding to NrCAM or β2-Spectrin by immunoprecipitation. Results revealed a pocket in ANK repeat R11 of the AnkB MBD that targets the NrCAM FIGQY motif. A maternally inherited ASD missense mutation in AnkB (A368G) located in this pocket disrupted association with NrCAM and functionally impaired Sema3F-induced spine pruning in cortical neurons. The AnkB/β2-Spectrin model identified key interactions between the AnkB SBD and spectrin repeats 14 and 15. A maternally inherited ASD missense mutation in the Zu51 (R977Q) subdomain of the AnkB SBD disrupted association with β2-Spectrin. These new findings provide insight into the L1-CAM/AnkB complex and molecular basis of ASD etiology associated with AnkB missense mutations.

## Results

### Structural Modeling and Experimental Validation of the AnkB-NrCAM Complex

Our initial goal was to predict key amino acids mediating binding between AnkB and the L1-CAM cytoplasmic domain, and then to assess whether ASD variants in the predicted AnkB region might alter the interactions. The lack of experimentally resolved structures or appropriate templates for homology modeling posed significant challenges for accurate structure prediction. To address this, we employed the deep learning-based protein structure prediction tool – AlphaFold-v2.2. We focused on NrCAM due to its role in Sema3F-induced spine pruning mediated by AnkB-220 (30). The cytoplasmic domain of NrCAM (residues 1191–1304, UniProt ID: Q92823) was modeled in association with ANK repeats 1–24 of the MBD (residues 30–822, UniProt ID: Q01484). The structure yielded a predicted TM-score (pTM) of 0.73 and inter-chain predicted TM score (ipTM) of 0.42, indicating moderate confidence in the overall fold of the complex (Fig. 2A).

**Figure 2:**
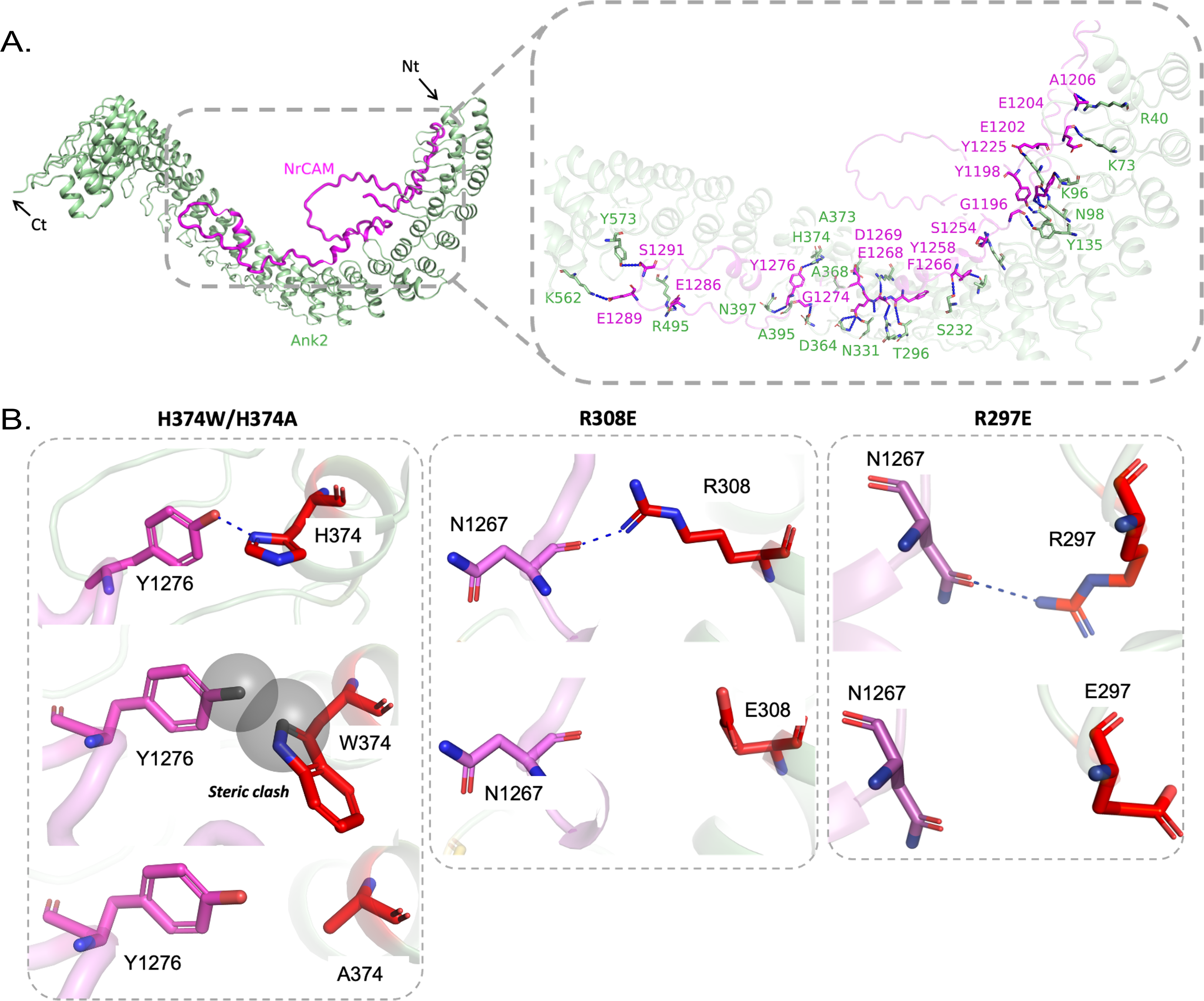
AlphaFold Model of the AnkB Membrane-Binding Domain Bound to the NrCAM Cytoplasmic Domain A. AlphaFold-predicted structure of the AnkB/NrCAM complex, showing the NrCAM cytoplasmic domain (residues 1191–1304, magenta) bound to the AnkB membrane-binding domain (MBD), comprising ANK repeats 1–24 (green). Arrows indicate the N-and C-terminal ends (Nt, Ct). B. Structural models generated using the PyMOL mutagenesis tool illustrate how AnkB mutations H374W, H374A, R308E, and R297E (red) alter interactions with NrCAM residues Y1276 and N1267 (magenta).

The AlphaFold model of the NrCAM-AnkB complex revealed critical interactions between the cytoplasmic domain of NrCAM and AnkB (Fig. 2A). Interaction analysis by PDBSum identified 8 salt bridges,18 hydrogen bonds, and 415 non-bonded contacts between AnkB and NrCAM (Fig. S1). A disordered loop in the NrCAM sequence of about 40 amino acids is seen protruding from the helical ANK repeats of AnkB. In the model Y1276 located in the FIGQY motif of NrCAM, formed a strong hydrogen bond with H374 of AnkB in ANK repeat R11, potentially stabilizing the complex (Fig. 2B, Fig. S1). In addition to hydrogen bonds, hydrophobic contacts were identified between surrounding residues, reinforcing the interaction interface. The NrCAM/AnkB interaction site constituted a pocket that also contained two residues AnkB A368 (55) and AnkB A373 (47) (Fig. 2) that are sites of genetic variation in ASD.

To test the model, mutations were generated in the 220 kD isoform of AnkB by site-directed mutagenesis of the cDNA sequence. Because AnkB-220, not AnkB-440, mediates Sema3F-Fc induced spine pruning in neuronal cultures (30), all mutations were engineered into AnkB-220 cDNA with a C-terminal 2X HA-tag. Mutations were confirmed by whole plasmid DNA sequencing.

Expression plasmids encoding wild type (WT) AnkB-220-HA and mutant AnkB-220-HA proteins were co-transfected with plasmids encoding WT NrCAM cDNA in equimolar amounts into HEK293 cells, and assayed for association by co-immunoprecipitation from cell lysates prepared in the nonionic detergent Brij98. AnkB-HA was immunoprecipitated with anti-HA antibodies adsorbed to Protein A/G magnetic beads. Immune complexes were separated by SDS-PAGE and transferred to nitrocellulose. Western blotting was carried out using NrCAM antibodies, followed by reprobing for AnkB with anti-HA antibodies.

NrCAM effectively co-immunoprecipitated with WT AnkB, as seen from associated NrCAM proteins of 220 kDa and ∼130 kDa in a representative blot (Fig. 3A). Nonimmune IgG did not pull down AnkB or NrCAM (Fig. 3A). We compared co-immunoprecipitation of NrCAM with WT AnkB vs. AnkB containing Trp or Ala substitutions at the H374 residue, as these mutations were predicted to perturb the interaction. Relative levels of NrCAM (220-and ∼130 kDa combined) and AnkB-HA were quantified by densitometric scanning of labeled bands (Fig. 3A). AnkB H374W and H374A were found to be strongly inhibited for NrCAM association. Quantification of blots from multiple replicate experiments (Fig. S2) yielded mean NrCAM/AnkB ratios (± SEM) that were significantly decreased for both AnkB mutants relative to WT (Fig. 3A; two-tailed t-test, *p < 0.05). NrCAM and other L1-CAMs (200 kDa) are proteolytically processed to N-terminal 130-150 kDa and 60 kDa C-terminal fragments (56,57). The fragments remain attached to the cell by homophilic and heterophilic binding interactions (58,59), although some may be released. They are the predominant species in mouse brain and vary in abundance depending on cell type, environment, and signaling (60). NrCAM antibodies to the extracellular domain were used here that recognize only the 220 kDa and ∼130 kDa proteins.

**Figure 3:**
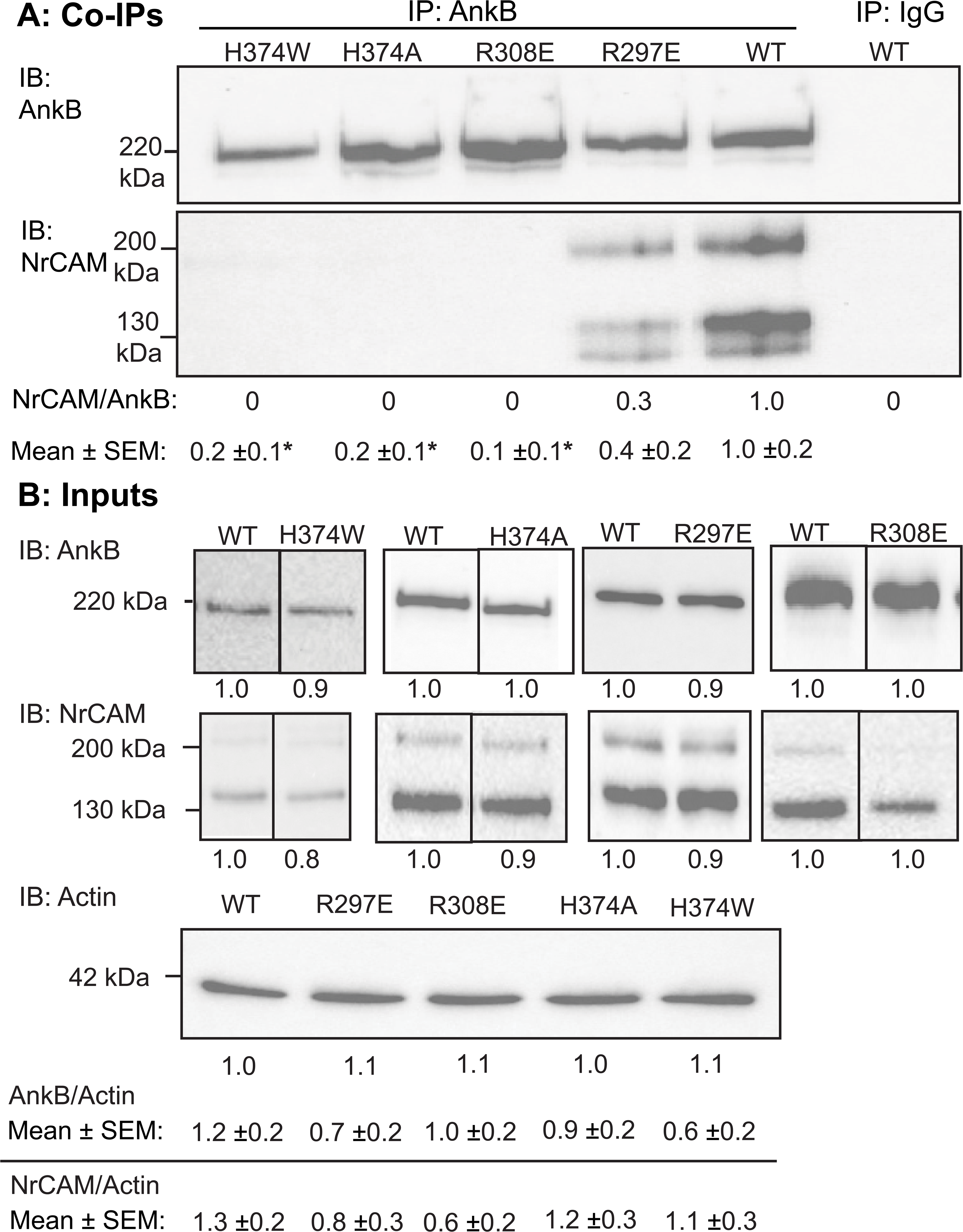
AnkB Associates with NrCAM at a Nexus of Residues in ANK Repeat R11 A. NrCAM co-immunoprecipitates with WT AnkB-220 (HA-tagged) from transfected HEK293 cells, and to a much lesser extent with HA-tagged AnkB-220 bearing point mutations H374W, H374A, R308E, and R297E. AnkB was immunoprecipitated (IP) from cell lysates with anti-HA antibodies, and immunoblotted (IB) with anti-NrCAM antibodies followed by reprobing with anti-HA antibodies. NrCAM/AnkB ratios in the immunoprecipitates relative to WT are shown under each lane of a representative blot. Mean ratios of NrCAM/AnkB ± SEM from 5 replicate experiments for each WT or mutant are indicated below. Significant mean differences (2-tailed t-test, p< 0.05) between WT and mutant AnkB were: * H374W (p= 0.002), * H374A (p=0.003), * R308E (p< 0.001), and * R297E (p=0.03). Nonimmune IgG did not immunoprecipitate AnkB or NrCAM. B. Input WT and mutant HA-AnkB-220 levels in HEK293 lysates (5 µg) prior to immunoprecipitation were determined by immunoblotting (IB) with anti-HA antibodies, followed by stripping and reprobing with anti-NrCAM antibodies, and then with anti-Actin antibodies as loading control. Mutant values relative to WT in each representative blot are shown below the lanes. Lanes from the same blot are shown without a line, or with a line if lanes were not adjacent. Blots from different gels are shown with a wider separation. Mean ratios of AnkB/Actin or NrCAM/Actin ± SEM from n replicate experiments are indicated below. There was not a significant mean difference in AnkB/Actin (2-tailed t-test, p > 0.05) between WT AnkB (n=12) and H374W (n=5, p= 0.08), H374A (n=6, p=0.31), R297E (n=5, p=0.13), R308E (n=6, p=0.08). There was also not a significant mean difference in NrCAM/Actin between WT and H374W (p= 0.39), H374A (p=0.54), R297E (p=0.09), and R308E (p=0.06). The position of the 220 kDa molecular weight marker from PageRuler Plus is indicated and coincides with the AnkB220-HA band.

To better understand their impact on the complexes, the PyMOL mutagenesis tool was employed to model the mutations. Substitution of H374 with tryptophan in the AnkB structural model introduced steric clashes with Y1276 in the FIGQY motif due to the larger side chain of tryptophan, disrupting the H-bond and causing loss of structural integrity within the AnkB-NrCAM complex (Fig. 2B).

Furthermore, replacing H374 with alanine prevented formation of the crucial H-bond with Y1276, reducing the stability of the complex and weakening the overall AnkB-NrCAM interface (Fig. 2B).

The PyMOL mutagenesis tool also predicted that AnkB R308 would promote the stability of the AnkB/NrCAM complex by forming a stabilizing H-bond with N1267 in the NrCAM cytoplasmic domain sequence QFN^1267^EDGSFIGQY^1276^(Fig. 2B). A charge substitution replacing R308 with glutamate (R308E) could prevent this interaction, destabilizing the local conformation. We generated the R308E substitution in AnkB-220 and assayed its association with NrCAM by co-immunoprecipitation from transfected HEK293 cells. This mutation resulted in nearly a complete loss of binding to NrCAM (Fig. 3A, Fig. S2). Additionally, N1267 in NrCAM potentially forms a stabilizing salt bridge with R297 in AnkB (Fig. 2B). Substitution of Glu for Arg at position 297 could disrupt this interaction, abolishing the salt bridge. Accordingly, the charge substitution R297E in AnkB was generated and found to decrease co-immunoprecipitation with NrCAM by ∼60% (Fig. 3A, Fig. S2).

To evaluate whether the AnkB mutant proteins were stably expressed and not degraded, HEK293 cells were transfected with WT or mutant AnkB cDNAs, together with NrCAM cDNA in equimolar quantities. Equal amounts of cell lysate protein were analyzed by Western blotting. AnkB and NrCAM protein bands were quantified by densitometric scanning and normalized to corresponding WT levels. As shown in representative blots, AnkB mutants H374W, H374A, R297E, and R308E were expressed at equivalent levels to WT, as was NrCAM (Fig. 3B). Blots were reprobed for endogenous Actin as a loading control. Evaluating AnkB/Actin and NrCAM/Actin ratios, all AnkB mutants and NrCAM were expressed at levels comparable to WT (Fig. 3B). Replicate experiments yielded similar results (Fig. S2). Mean AnkB/Actin and NrCAM/Actin ratios from replicate assays are presented below in Fig. 3B.

### ASD-Related Variants in AnkB are Impaired for NrCAM Binding

We next assessed interactions of human ASD missense variants selected in the AnkB MBD that were predicted by AlphaFold-v2.2 to impact NrCAM binding. We first analyzed AnkB A368G, which is a maternally inherited missense mutation in ANK repeat R11 (47,55) The mutation was generated by site-directed mutagenesis in AnkB-220 cDNA, and analyzed for NrCAM binding by co-immunoprecipitation from transfected HEK293 cell lysates. AnkB A368G was significantly impaired for binding to NrCAM, as evident from the decreased NrCAM/AnkB ratio compared to WT in the immunoprecipitates on a representative blot (Fig. 4A). Quantification of replicate blots (Fig. S3) yielded mean NrCAM/AnkB ratios (± SEM) that were significantly decreased for AnkB A368G compared to WT (Fig. 4A; two-tailed t-test, *p < 0.05). The substitution of alanine with glycine at position 368 likely increases local conformational flexibility due to glycine’s smaller side chain and lack of β-carbon, destabilizing the binding pocket for NrCAM. Additionally, the AnkB R297-NrCAM N1267 salt bridge may be affected by increased flexibility in the A368G variant, further influencing optimal NrCAM binding.

**Figure 4:**
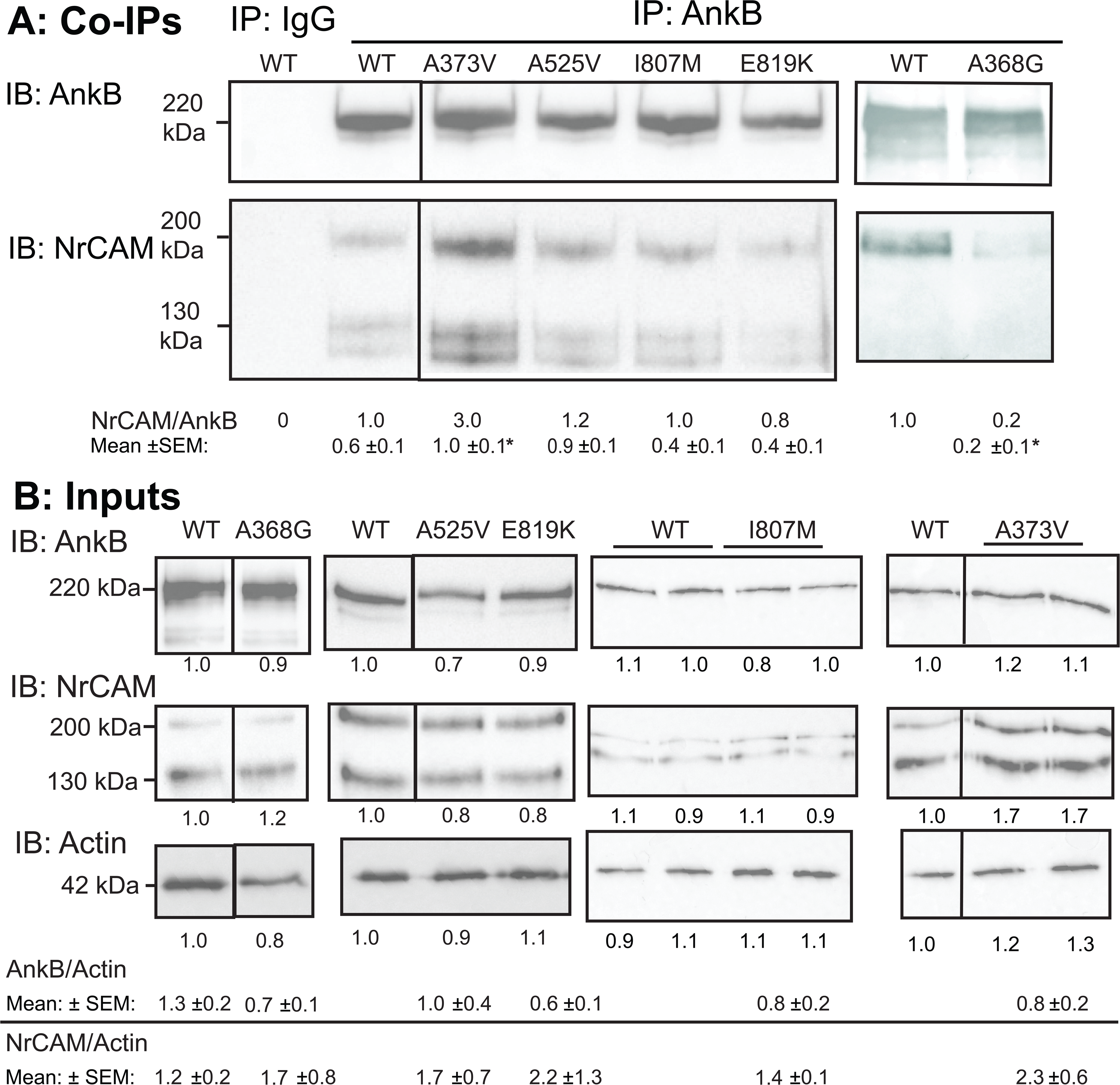
Altered Interactions of Missense ASD Mutations in AnkB A368G and A373V with NrCAM A. NrCAM co-immunoprecipitated (co-IP) with HA-tagged WT AnkB-220 from transfected HEK293 cells with anti-HA antibodies but not nonimmune IgG. ASD mutation A373V showed enhanced association with NrCAM; A368G mutation showed decreased association with NrCAM. NrCAM/AnkB ratios were obtained by densitometry as presented under the representative blots. Lanes from the same blot are shown without a line or with a line if not adjacent. Blots from different gels are shown with a wider separation. Mean ratios of NrCAM/AnkB ± SEM from n replicate experiments for WT and mutants are indicated below. Mean differences (2-tailed t-test, *p< 0.05) were significant between WT and mutant AnkB for *A373V (n=3, p= 0.02) and * A525V (n=3, p=0.15), but not for I807M (n=5, p=0.15) or E819K (n=3, p=0.23). A. Input WT and mutant HA-AnkB-220 levels in HEK293 lysates (5 µg) prior to immunoprecipitation were determined by immunoblotting (IB) with anti-HA antibodies, followed by stripping and reprobing with anti-NrCAM antibodies, then with anti-Actin antibodies as loading control. Lanes from the same blot are shown without a line or with a line if not adjacent. Blots from different gels are shown with a wider separation. Levels of AnkB, NrCAM, and Actin relative to WT are shown below representative blots. Mean ratios of AnkB/Actin ± SEM from n replicate experiments indicated that there was not a significant difference in AnkB/Actin (2-tailed t-test, p< 0.05) between WT (n=10) and mutant AnkB for A368G (n=3, p=0.35), A525V (n=3, p=0.68), E819K (n=4, p=0.27), I807M (n=4, p=0.15), or A373V (n=4, p= 0.40). There was also not a significant mean difference in NrCAM/Actin between WT and A368G (p= 0.43), A525V (p=0.77), E819K (p=0.40), I807M (p=0.39), or A373V (p=0.43). The position of the 220 kDa molecular weight marker from PageRuler Plus is indicated and coincides with the AnkB220-HA band.

Other maternally inherited ASD mutations A373V in R11 (47), A525V in R15 (55), and E819K in R24 (55) were similarly assayed, along with a *de novo* AnkB mutation I807M in R24 (47,61). AnkB Surprisingly, the neutral A373V mutation increased binding to NrCAM (Fig. 4A). This might be explained by the bulkier isopropyl group of the valine side chain than the methyl group of the alanine side chain, which could induce additional hydrophobic interactions to increase stability. In contrast, AnkB variants A525V, I807M, and E819K were not significantly impacted for binding to NrCAM (Fig. 4A). Based on PDBSum the neutral mutations A525V and I807M were not expected to alter key interactions nor introduce new ones (Fig. S1), suggesting a minimal impact on the AnkB-NrCAM complex’s overall stability. Structural modeling predicted that despite its being a charge variant AnkB E819K was unlikely to have a binding interface with NrCAM. For these ASD mutants, replicate experiments gave similar results (Fig. S3), and quantification yielded mean NrCAM/AnkB ratios (± SEM) that were not significantly decreased relative to WT (Fig. 4A; two-tailed t-test, p > 0.05).

To evaluate whether the ASD-related AnkB mutant proteins were stably expressed, lysates of HEK293 cells were transfected with WT or mutant AnkB, together with equimolar amounts of NrCAM plasmid, and equal amounts of protein were analyzed by Western blotting. All of the mutants were expressed at equivalent levels to WT AnkB with similar AnkB/Actin ratios (Fig. 4B). Replicate experiments yielded similar results (Fig. S4) with mean NrCAM/AnkB ratios (± SEM) that were not significantly decreased relative to WT (Fig. 4B; two-tailed t-test, p > 0.05).

### Structural Modeling and Experimental Validation of the AnkB-β2 Spectrin Complex and Interaction with ASD Variants

Spectrin β1-4 proteins are present in spines and dendrites of pyramidal neurons, and β2-Spectrin (*SPTBN1*; UniProt ID: Q01082) is the most robustly expressed at these sites (62–65). The AnkB/β*2-*Spectrin complex was modeled using the AlphaFold-v2.2 approach to identify key molecular interactions and evaluate potential impact of ASD mutations on its structure. Given that the SBD of AnkB engages β2-Spectrin at repeats 14 and 15 (66), we focused our analysis on the interface between AnkB residues 966–1125 within the SBD, and β*2-*Spectrin residues 1563–2093 encompassing spectrin repeats 14 and 15. The predicted structure yielded a pTM score of 0.71 and ipTM score of 0.69, indicating high confidence in the overall fold of the complex (Fig. 5A).The AlphaFold model of the AnkB/β*2-*Spectrin complex revealed important interactions between these regions (Fig. 5). Residue-level interaction analysis identified 2 salt bridges, 10 H-bonds, and 145 non-bonded contacts between AnkB and β*2-*Spectrin interfaces (Fig. S4).

**Figure 5:**
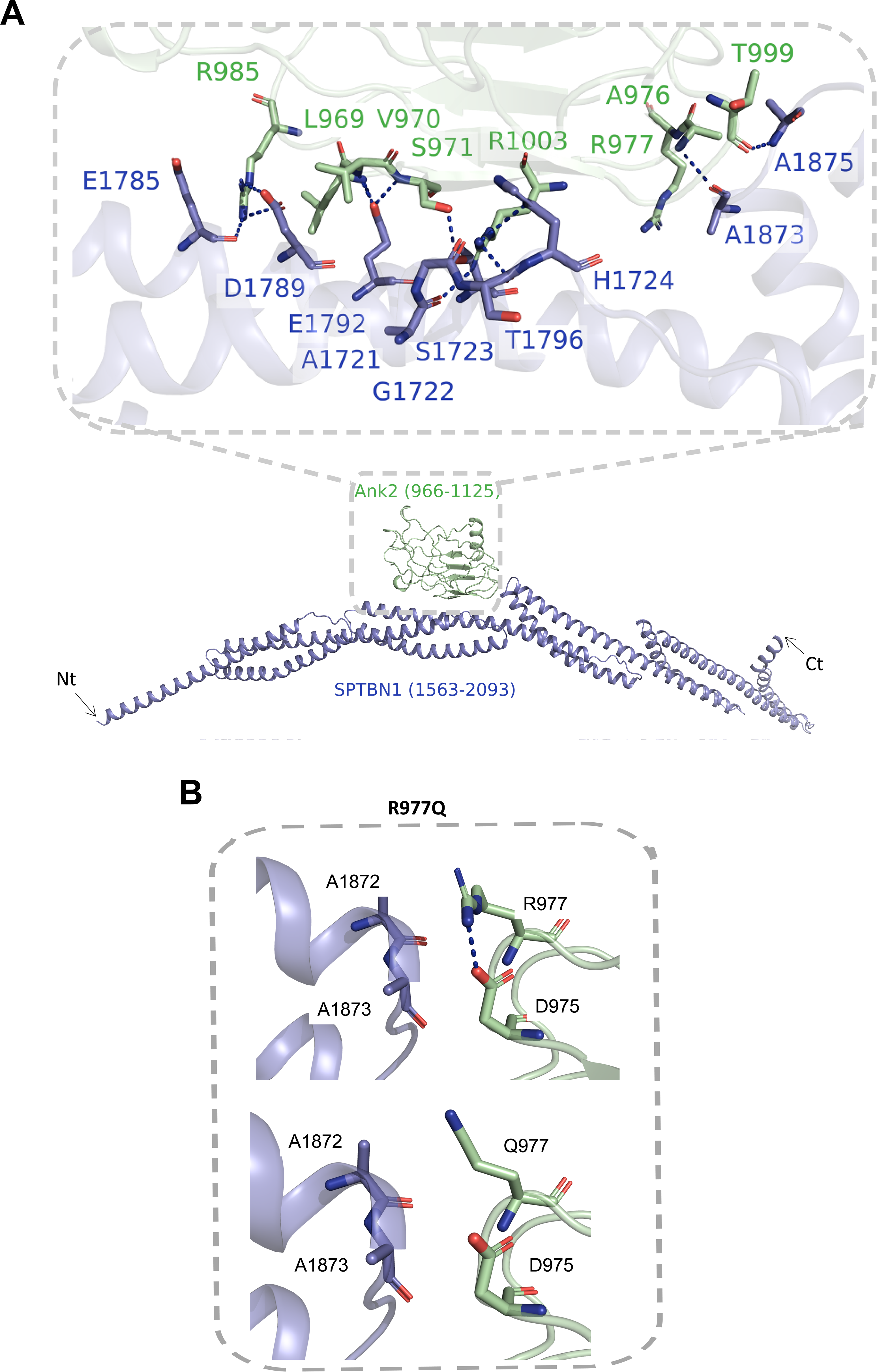
AlphaFold Model of the AnkB Spectrin-Binding Domain Interaction with β2-Spectrin (SPTBN1) A. AlphaFold-predicted structure of the AnkB/ β2-Spectrin complex, showing the AnkB spectrin-binding domain (residues 966–1125, green) bound to spectrin repeats 14 and 15 of β2-Spectrin (residues 1563–2093, blue). Arrows indicate the N-and C-terminal ends (Nt, Ct). Dashed blue lines represent hydrogen bonds. B. Structural model generated using the PyMOL mutagenesis tool illustrating an intramolecular interaction between AnkB (green) residues R977 and D975, and β2-Spectrin (blue) at residues A1872 and A1873. The ASD-associated AnkB 977Q mutation disrupts this intramolecular interaction, potentially weakening its binding to β2-Spectrin. The dashed blue line represents a salt bridge.

We next assessed protein-protein interactions of two ASD missense mutations predicted to impact β2-Spectrin binding in the AnkB SBD: a maternally inherited mutation R977Q in the ZU5A region (4) and *de novo* mutation P1380R in the UPA region (47). We generated the two mutations in HA-tagged AnkB-220 cDNA and co-expressed them with β2-Spectrin-GFP cDNA in HEK293 cells. β2-Spectrin-EGFP was pulled down from cell lysates with anti-GFP antibodies adsorbed to Protein A/G magnetic beads. Immune complexes were analyzed by Western blotting with anti-HA antibodies, followed by re-probing with anti-GFP antibodies. Relative levels of β2-Spectrin/AnkB were quantified by densitometry. AnkB-220 R977Q was impaired for binding to β2-Spectrin, as evident from the decreased AnkB/β2-Spectrin-GFP ratio in co-immunoprecipitates relative to WT (Fig. 6A).

**Figure 6:**
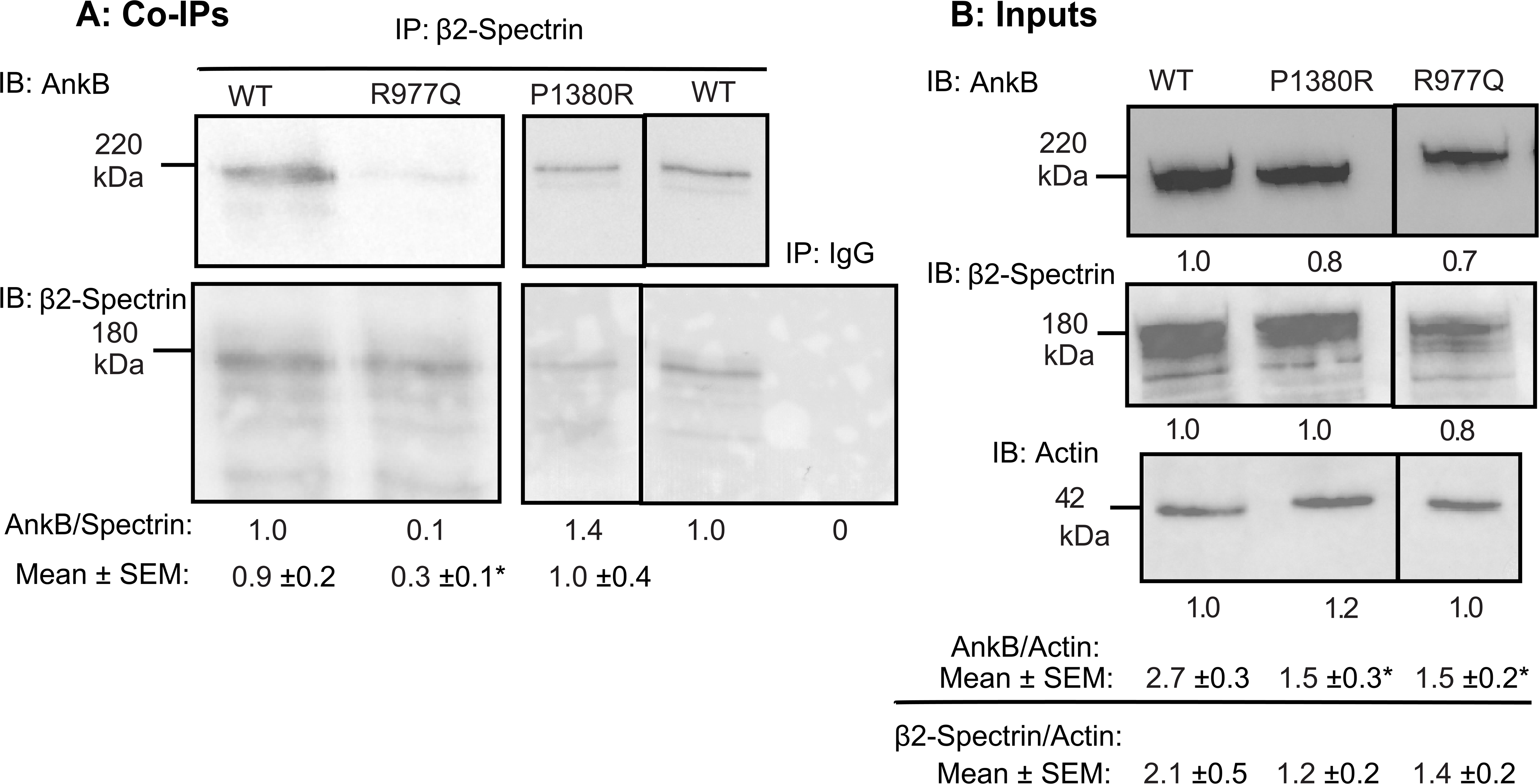
Altered Interaction of ASD Missense Mutation in AnkB R977Q with β2-Spectrin A. AnkB-220 was co-immunoprecipitated with GFP-tagged β2-Spectrin from transfected HEK293 cells using anti-GFP antibodies. Inherited ASD mutation AnkB R977Q showed significantly decreased association with β2-Spectrin, but *de novo* mutation AnkB P1380R was not impaired for binding. AnkB/β2-Spectrin ratios in the immunoprecipitates relative to WT are shown under the representative blots. Lanes from the same blot are shown without a line, or with a line if not adjacent. Blots from different gels are shown with a wider separation. Mean ratios of AnkB/β2-Spectrin ± SEM from n replicate experiments are indicated below. Mean differences (2-tailed t-test, *p< 0.05) were significant between WT (n=12) and * R977Q (n=7, p= 0.02) but not for P1380R (n=6, p=0.82). B. Input WT and mutant AnkB-220 levels in HEK293 lysates (5 µg) prior to immunoprecipitation were determined by immunoblotting (IB) with anti-HA antibodies, followed by stripping and reprobing with anti-EGFP antibodies, and anti-Actin antibodies as loading controls. Lanes from the same blot are shown without a line, or with a line if not adjacent. Levels of AnkB, β2-Spectrin, and Actin relative to WT in the representative blots are shown below each lane. Mean ratios of AnkB/Actin ± SEM and β2-Spectrin/Actin from n replicate experiments are indicated below. There was a significant mean difference in AnkB/Actin (2-tailed t-test, *p< 0.05) between WT (n=4) and mutants R977Q (n=4, p= 0.02) and P1380R (n=4, p=0.02).There was not a significant mean difference in β2-Spectrin/Actin between WT and R977Q (p=0.31) or P1380R (p=0.17). The position of the 220 kDa molecular weight marker from PageRuler Plus is indicated and coincides with the AnkB220-HA band.

Quantification of replicate blots (Fig. S5) yielded mean NrCAM/AnkB ratios (± SEM) that were significantly decreased for AnkB R977Q compared to WT (Fig. 6A; two-tailed t-test, *p < 0.05). Computational prediction of this disruption suggested that the mutation disrupted a salt-bridge interaction between AnkB R977 and D975, and weakened hydrophobic interactions with β2-Spectrin A1872 and A1873 (Fig. 5B). Thus, the prediction was consistent with experimental mutagenesis data showing reduced AnkB/β2-Spectrin binding. In contrast, AnkB-220 P1380R did not exhibit decreased affinity for β2-Spectrin in the co-immunoprecipitation assay, as evident from mean AnkB/β2-Spectrin ratios in immunoprecipitates in replicate assays (Fig. 6A, Fig. S5). The P1380R mutation is located C-terminally in a disordered region outside the AlphaFold modeled region of the AnkB SBD.

To evaluate whether these AnkB mutant proteins were stably expressed, lysates of HEK293 cells were transfected with WT or mutant AnkB plasmids, together with equimolar levels of β2-Spectrin-GFP cDNA, and equal amounts of cell lysate protein were analyzed by Western blotting. The two ASD-related AnkB mutants were expressed at reduced levels compared to WT AnkB relative to Actin as shown in the representative blot (Fig. 6B). Replicate experiments yielded similar results (Fig. S5) with mean AnkB/Actin or AnkB/Vinculin ratios (± SEM) that were approximately 60% of WT (Fig. 6B; two-tailed t-test, *p < 0.05). β2-Spectrin levels were similar in mutant vs. WT expressing cells relative to Actin (Figs. 6B, S5). These results suggested that the AnkB R977Q and P1380R proteins are partly unstable in HEK293 cells.

### Sema3F-induced Spine Retraction is Promoted by WT AnkB-220 but not by AnkB-A368G

Studies in transgenic mouse models and neuronal cultures have demonstrated that Sema3s trigger spine retraction on apical dendrites through NrCAM, Neuropilin-2, and AnkB-220 (21–23,30,67). Because AnkB A368G was the most compromised ASD missense mutation for NrCAM interaction, we assessed whether it was deficient in spine pruning using an established assay for Sema3F-induced dendritic spine retraction in cultured cortical neurons from Ank2 mutant mice.

Neurons from *Ank2^+/-^* heterozygotes and *Ank2^-/-^* homozygotes have been shown to be inhibited for Sema3F-induced spine retraction in this *in vitro* assay (30). Cortical neuronal cultures from *Ank2^+/-^* heterozygous embryos (E15.5) were co-transfected on DIV11 with AnkB-220 WT or mutant cDNA together with pCAG-IRES-mEGFP plasmids. Neurons in replicate cultures were treated on DIV14 with Sema3F-Fc or control Fc (5 nM) for 30 min and spine density quantified on apical dendrites in deconvolved confocal z-stacks using Neurolucida software. Mean spine density of neurons expressing WT AnkB-220 or mutant A368G AnkB-220 were compared by the t-test (two-tailed, p < 0.05). Neurons transfected with WT AnkB-220 responded to Sema3F-Fc with spine retraction that was significantly greater than that of control Fc treated neurons (Fig. 7). In contrast, neurons expressing A368G AnkB-220 did not respond to Sema3F-Fc with significant spine retraction (Fig. 7). Representative images of apical dendrites from WT or mutant transfected cultures treated with Fc or Sema3F-Fc are shown in 3D reconstructions performed in Imaris (Fig. 7), and as raw confocal images (Fig. S6).

**Figure 7:**
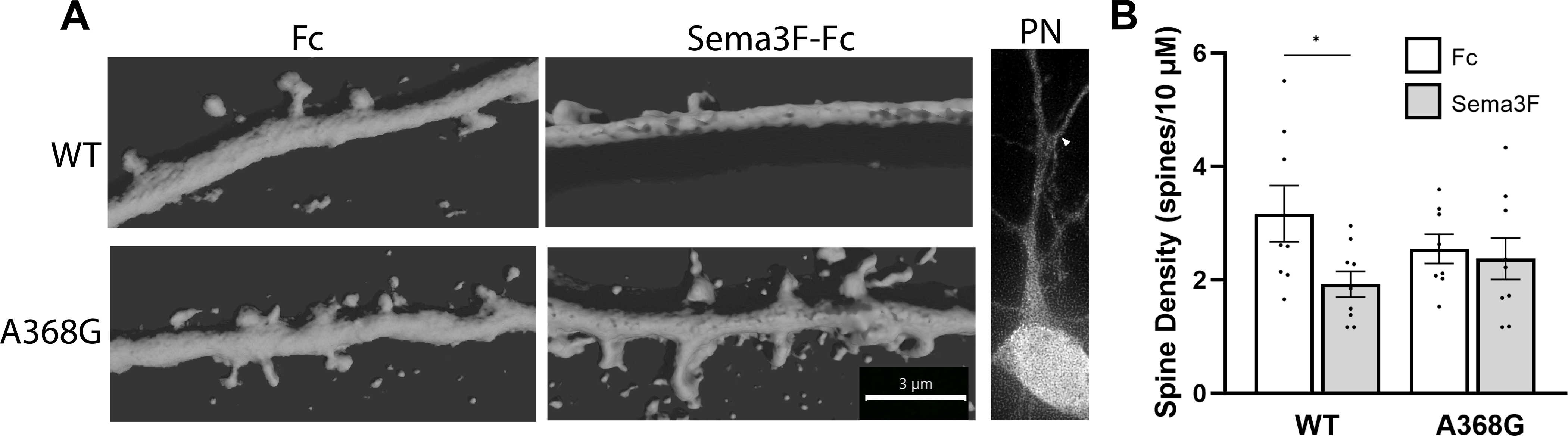
Sema3F-Induced Spine Retraction is Impaired in Neuronal Cultures from *Ank2 ^+/-^* Mice Expressing the ASD Mutation AnkB A368G. A. *Ank2 ^+/-^*cortical neuronal cultures were transfected with pCAG-IRES-EGFP together with plasmids expressing WT AnkB-220 or AnkB-220 mutant A368G. Cells were treated on DIV14 with 5 nm Fc (control) or Sema3F-Fc for 30 min, and immunostained for EGFP. Representative confocal images of EGFP-labeled apical dendrites on pyramidal neurons were subjected to 3D reconstruction in Imaris. Scale bar = 3 µm, all panels. Low magnification confocal image of pyramidal neuron (PN) with arrowhead pointing to the first branch of the apical dendrite (A). B. Quantification of mean spine density ± SEM per neuron on apical dendrites in neuronal cultures described in A. Each point presents the mean spine density per neuron per 10 µm length of apical dendrite. Results show that Sema3F-Fc induces significant spine pruning relative to Fc controls in *Ank2^+/-^* cortical neurons expressing WT AnkB but not ASD mutant AnkB A368G. P-values for Sema3F-Fc vs Fc (two tailed t-test, unequal variance) were for WT (* p= 0.04) and A368G (p=0.71). n=123-172 pines on 10 pyramidal neurons per condition.

## Discussion

AnkB, encoded by the high confidence ASD gene *Ank2*, regulates dendritic spines and excitatory synapses of cortical pyramidal neurons, and mediates Sema3F-induced spine pruning through NrCAM (30). To better understand the role of AnkB in these neuronal functions, molecular modeling of the AnkB MBD in complex with the NrCAM cytoplasmic domain and β2-Spectrin was undertaken through the use of AlphaFold. The predicted interactions were tested by substitution mutagenesis of the *Ank2* sequence and protein-protein co-immunoprecipitation from transfected HEK293 cells. This approach defined a binding pocket in the R11 ANK repeat of AnkB for the NrCAM cytoplasmic sequence QFN^1267^E^1268^DGSFIGQY^1276^, which is highly conserved in the L1-CAM family.

Critical residues R297, R308, and H374 in the AnkB MBD were identified that engaged NrCAM residues E1268, N1267, and Y1276, respectively, at the binding interface. Substitution mutagenesis of these AnkB residues resulted in significant losses of NrCAM binding. Alphafold modeling predicted that the inherited ASD mutation AnkB A368G is within an α-helical segment of the MBD near the NrCAM binding interface. The AnkB A368G mutation markedly disrupted association with NrCAM, despite being stably expressed. The ASD missense mutation AnkB A373V, located within the NrCAM binding interface adjacent to H374, significantly increased binding to NrCAM. Increased stability could influence molecular dynamics of the AnkB/NrCAM complex or phosphorylation of Y1276 in the FIGQY motif, which promotes Ankyrin dissociation from L1-CAMs (34). Other AnkB ASD missense mutations at or near the MBD-A525V, I807M, and E819K-were stably expressed but had little effect on NrCAM binding. These variants were not predicted in our models to exhibit a binding interface with NrCAM. ASD pathology associated with these *Ank2* mutations could be due to altered interactions with other binding partners, or to other genes.

AlphaFold modeling of the AnkB/β2-Spectrin complex identified key interactions between the AnkB SBD and β2-Spectrin repeats 14 and 15. β2-Spectrin (SPTBN1) variants have been linked to ASD and related syndromes (45,53,54). The ASD missense mutation (R977Q) in the Zu51 subdomain of the AnkB SBD was found to disrupt association with β2-Spectrin. In contrast, the ASD missense mutation P1380R was not impaired for β2-Spectrin binding. Both AnkB R977Q and P1380 exhibited reduced levels of expression in HEK293 cells. This decrease in stability might impact known β2-Spectrin functions in dendrite development, cell membrane organization, or transport (reviewed in (45)). However, the stability of these mutants in neurons or consequences on neuronal function has not been evaluated.

The AnkB A368G mutation markedly disrupted association with NrCAM and impaired Sema3F-induced spine retraction in mouse cortical neuron cultures. Previous structure-function studies demonstrated that binding between the extracellular domains of NrCAM and Npn-2 increases the affinity of Npn-2 for PlexA3 to activate Sema3F-induced signaling through Rap-GAP and spine pruning (24). AnkB binding to the NrCAM cytoplasmic domain may further stabilize the Sema3F holoreceptor NrCAM/Npn-2/PlexA3 at the neuronal membrane through linkage to the underlying actin cytoskeleton. Sema3F also induces Npn-2 endocytosis together with GluA1 AMPA receptors, decreasing firing during homeostatic downscaling (68). Although consequences of AnkB A368G *in vivo* have not been determined, loss of NrCAM/AnkB binding could, in principle, contribute to the ASD phenotype by inhibiting developmental spine pruning, consistent with increased spine density in frontal brain regions in ASD cases (14–18).

It is not known if binding of NrCAM to AnkB alters the AnkB/Nav1.2 interaction at the pyramidal cell dendritic membrane. Nav1.2 is important for back-propagation of action potentials in dendrites, which could impair synaptic function and plasticity (29). Although the binding sites for Nav1.2 and the L1-CAM Neurofascin in the AnkB MBD partially overlap, they have not been mapped precisely (44).

Thus, it will be interesting to compare AlphaFold models of AnkB with NrCAM and Nav1.2, and to determine if AnkB A368G or other MBD mutations affect Nav1.2 binding. A limitation of the AlphaFold approach is that neither the structure of disordered regions in AnkB, β2-Spectrin, and L1-CAMs, nor alternate conformations of AnkB (active/inactive) could be modeled.

AnkB ASD mutations might alter other critical neuronal functions carried out by AnkB and L1-CAMs, such as synaptic stabilization. Both AnkB and NrCAM are required for stabilization of perisomatic synaptic contacts made by cholecystokinin-expressing basket interneurons onto cortical pyramidal cells (69). Ankyrin also stabilizes GABAergic inputs to pyramidal cell soma through L1, as demonstrated in mice with a FIGQY to H mutation in L1 (39). Additionally, L1 binding to related Ankyrin G drives chandelier interneuron innervation of the axon initial segment of pyramidal neurons (40). A consideration is that all of the mutations studied here target both AnkB-220 and AnkB-440, and thus increased axonal branching and ectopic connectivity due to AnkB-440 deficiency could also be impacted (31,32). Future *in vivo* studies will be needed to assess functional consequences of AnkB ASD mutations. For example, new mouse mutational models can be generated and tested for neuronal pathology and behavioral phenotypes relevant to ASD. In summary, these new findings provide insight into the L1-CAM/AnkB/β2-Spectrin complex and molecular basis of ASD etiology associated with AnkB missense mutations.

## Experimental Procedures

### Structural Modeling of AnkB-NrCAM and AnkB-β2-Spectrin Complexes

The structural models of AnkB and its interactions with NrCAM and β2-Spectrin (SPTBN1) were predicted using AlphaFold v2.2 (70), a state-of-the-art deep learning algorithm for protein structure prediction. The cytoplasmic domain of human NrCAM (residues 1191–1304, UniProt ID: Q92823) was modeled in association with ANK repeats 1–24 in the human AnkB MBD (residues 30– 822, UniProt ID: Q01484). Similarly, the complex between AnkB human β2-Spectrin (UniProt ID: Q01082) was modeled, focusing on the interface between AnkB residues 966–1125 and β2-Spectrin residues 1563–2093. To enhance the accuracy of the predicted models, the complex structures were further optimized using the AMBER molecular dynamics package, employing energy minimization and equilibration protocols (71). Interactions between AnkB and NrCAM or β2-Spectrin (*SPTBN1*) were visualized using the PyMOL Molecular Graphics System, Version 3.0 Schrödinger, LLC and Chimera-X (72). The PDBsum tool (73) was utilized to identify detailed inter-protein interactions. Electrostatic interactions and the charged surface distributions of the complexes were analyzed using Advanced Poisson-Bolzmann Solver software (APBS) (74). This tool was used to compute the electrostatic potential across the protein surfaces, enabling the identification of charged regions critical for complex formation. The PyMOL mutagenesis tool was employed to model specific mutations in AnkB, NrCAM, and β2-Spectrin. These mutations were structurally analyzed to evaluate their impact on binding interfaces and overall complex stability.

### Plasmids and Mutagenesis

Plasmids used were: *Ank2* cDNA encoding the human AnkB isoform of 220 kDal molecular weight with a C-terminal 2x hemagglutinin (HA) tag in pEGFP-N1-ΔEGFP (Addgene), mouse NrCAM cDNA (NCBI nm_01127493; variant 3 in pCMV6-XL4 (21)), human β2-Spectrin with C-terminal EGFP tag in pEGFP-N1 (75). Mutations were engineered into AnkB-220 by site-directed mutagenesis using Q5 mutagenesis (New England Biolabs). Sequences were confirmed by whole plasmid sequencing at Plasmidsaurus, Inc.

### Immunoreagents

Antibodies used were directed against the following proteins: rabbit anti-NrCAM (Abcam #24344 or R&D Systems AF8538), chicken anti-EGFP (Abcam #13970), rabbit anti-GFP (Invitrogen A6455), mouse anti-AnkB (Invitrogen 33-3700), mouse anti-Actin (EMD Millipore Ab1501), mouse anti-Hemagglutinin (HA) (Life Technologies, 2-2.2.14). Non-immune rabbit IgG (NIgG), HRP-and AlexaFluor 488, 555, and 594-conjugated secondary antibodies were from Jackson Immunoresearch. *HEK293 Cell Culture, Transfection, and Immunoprecipitation*

HEK293 cells were grown in DMEM/ gentamicin/kanamycin/10% FBS in a humidified incubator with 5% CO_2_. Cells were seeded at 2 × 10^6^ cells/100mm dish and transfected when cells reached ∼ 2 x 10^7^ cells/dish. AnkB 220-2x HA and NrCAM plasmids were transfected in equimolar quantities using Lipofectamine 2000 in Opti-MEM as described (22). Media was changed to DMEM after 18 hr, and cells were lysed and collected 48 hr post-transfection. Cells were harvested in Brij98 lysis buffer (1% Brij98, 10 mM Tris-Cl pH 7.0, 150 mM NaCl, 1mM EDTA, 1mM EGTA, 10 mM NaF, protease inhibitors (SigmaAldrich).

For co-immunoprecipitation (co-IP) of NrCAM and AnkB-220-HA, lysates of transfected HEK293 cells (0.5 mg) were diluted in co-IP binding buffer (PBS, 0.1% TritonX-100, protease inhibitors) and precleared by incubating with Pierce Protein A/G magnetic beads (ThermoFisher Scientific) for 30 minutes at 4°C. Precleared lysates (duplicates or triplicates, 500 ug each) were then incubated with fresh Protein A/G magnetic beads that had been coated with mouse anti-HA antibody or control NIgG overnight at 4°C. Beads were collected using a magnetic stand and washed 4x in co-IP binding buffer. Immune complexes were eluted in SDS sample buffer by boiling for 2 min. Proteins were separated by SDS-PAGE (6%) and transferred to nitrocellulose membranes. Membranes were blocked in TBST (Tris buffered saline/0.1% Tween-20) containing 5% nonfat dried milk and incubated overnight with primary antibodies (1:1000), washed, and incubated with HRP-secondary antibodies (1:5000) for 1 hr. Blots were first probed with rabbit polyclonal antibody to NrCAM (R &D 8538) and HRP-conjugated donkey anti-rabbit IgG. All antibodies were diluted in 5% nonfat dry milk/TBST. Blots were developed using Western Bright ECL Substrate (Advansta). Images were captured with a BioRad ChemiDoc imager for times yielding a linear response of signal. Blots were then stripped and reprobed for AnkB using mouse anti-HA monoclonal antibody and HRP-conjugated goat anti-mouse IgG. Densitometry of bands on TIFF images was measured in FIJI (mean gray value, 8 bit).

Background was subtracted using the rolling ball method with radius set to 50 pixels. The ratio of AnkB to NrCAM was calculated from replicate experiments, and means ± SEM compared between WT and mutants by the t-test (two-tailed, unequal variance, p < 0.05). WT and mutant bands on the same representative blot were normalized to WT.

For co-immunoprecipitation of AnkB-220-HA and β2-Spectrin-GFP, lysates of HEK293 cells (0.5 mg) transfected with equimolar quantities of plasmids were diluted in co-IP binding buffer and precleared by incubating with Protein A/G magnetic beads for 30 minutes at 4°C. Beads were removed using the magnetic stand. Precleared lysates (duplicates or triplicates, 500 ug each) were incubated with rabbit-anti-EGFP antibody or control IgG for 4 hr at 4°C, then with Protein A/G magnetic beads. Beads were collected using the magnetic stand and washed 4x in co-IP binding buffer. Immune complexes were eluted in SDS sample buffer, separated on a 6% SDS-PAGE gel and transferred to nitrocellulose membranes. Membranes were immunoblotted with mouse anti-HA antibody and HRP-conjugated goat anti-mouse IgG, and imaged on the ChemiDoc. Membranes were stripped and reprobed for β2-Spectrin-GFP using rabbit anti-GFP and HRP-donkey anti-rabbit IgG. Densitometry of bands on TIFF images was measured and background subtracted, as described above. The ratio of AnkB to β2-Spectrin-GFP was determined from replicate experiments, and means ± SEM compared between WT and mutants by the t-test (p < 0.05). Bands on representative blots were normalized to WT in figures.

For assaying the relative levels of WT and mutant AnkB in input HEK293 cell lysates prior to co-immunoprecipitation, equal amounts of protein (5 µg) determined by BCA protein analysis were subject to SDS-PAGE, transferred to nitrocellulose, and immunoblotted with the indicated antibodies. Membranes were reprobed first with antibodies to NrCAM or β2-Spectrin-GFP, followed by anti-Actin antibodies as loading controls. Images were collected with the BioRad ChemiDoc imager for times producing a linear response of signal. Densitometric signals of bands on representative blots were normalized to WT in the figures. Ratios of WT and AnkB relative to Actin were calculated and means ± SEM were compared between WT and mutants by the t-test (2-tailed, p < 0.05).

### Mouse Cortical Neuron Cultures and Sema3F-induced Spine Retraction

We previously described the production of cortical neuron cultures from *Ank2*-null and heterozyous mice on a first generation Sv129/C57Bl hybrid background (30). Cortical neurons from homozygous *Ank2*^-/-^ and heterozygous *Ank2*^+/-^ hybrids exhibited decreased spine collapse in response to Sema3F-Fc, which could be rescued by transfection of cDNA encoding WT AnkB-220 (30). *Ank2* heterozygous mice on C57BL/6 (76) were crossed with WT Sv129 mice to produce *Ank2*-heterozygous F1 hybrids. Intercrossing these F1 hybrids yielded WT, *Ank2*^+/-^, and *Ank2*^-/-^ genotypes. Because *Ank2*-/-embryos were present in lower than Mendelian ratios, we focused on *Ank2*-heterozygous neurons. Mice were maintained according to policies of the University of North Carolina Institutional Animal Care and Use Committee (IACUC; AAALAC Institutional Number: #329; ID# 18-073, 21-039) in accordance with NIH guidelines.

Cortical neurons were plated onto Lab-Tek II chamber slides (1.5 x 10^6^ cells/well) coated with poly-D-lysine and laminin as described (22). Genotyping of embryos was performed on tail DNA after plating the cells. AraC was added at 5 days *in vitro* (DIV5) to limit the growth of glia and fibroblasts, and media was changed on DIV7. At DIV11, cells were transfected with pCAG-IRES-mEGFP with WT or mutant AnkB-220 plasmids using Lipofectamine 2000 (22,24). At DIV 14, replicate cultures were treated with purified Fc from human IgG (Abcam #ab90285) or recombinant Sema3F-Fc fusion protein (R&D Systems #3237-S3) at 5 nM for 30 minutes. Cultures were fixed with 4% paraformaldehyde (PFA), quenched with 0.1M glycine, permeabilized with 0.1% Triton X-100, and blocked with 10% donkey serum. Cells were incubated with chicken anti-GFP primary antibody and AlexaFluor AF488-conjugated goat anti-chicken secondary antibody (1:500), washed, and mounted. At least 10 images of apical dendrites of labeled pyramidal neurons were captured per condition using a Zeiss Axioplan epifluorescence microscope equipped with a 40X oil objective. Images were deconvolved for scoring spine density on apical dendrites. Apical dendrites were differentiated from basal dendrites by cell morphology. Pyramidal neurons extend a single prominent, thicker apical dendrite from the apex of the pyramidal-shaped soma, whereas they extend multiple thinner basal dendrites from the base of the soma (77). Any ambiguous dendrites were not analyzed. Spines were scored blind to observer using Neurolucida software as described (30). Mean spine densities (number/10 µm ± SEM) were calculated and compared for significant differences by the t-test (2-tailed, unequal variance, p<0.05). In some instances confocal z-stacks were captured on a Zeiss LSM900 microscope using a Plan-apochromat 63 x 1.4 numerical aperture objective and Zen software in 0.2 µm optical sections. 3D reconstructions were rendered from dendritic z-stacks with Imaris (Bitplane) software. All experiments were designed to provide sufficient power (80–90%) to discriminate significant differences (p<0.05) in means (± SEM) between independent controls and experimental samples as described (78). The type I error probability associated with tests of the null hypothesis was set at 0.05.

## Data Availability

All data are contained within the article.

## Supporting Information

This article contains supporting information.

## Abbreviations

L1-CAMs: L1 cell adhesion molecules
NrCAM: neuron-glial related cell adhesion molecule
Npn-1/2: Neuropilin-1/2
PlexA: Plexin A
AnkB: Ankyrin B
*Ank2*: Ankyrin 2 gene
HA: hemagglutinin
*SPTNB1*: β2-Spectrin gene
ANK: ANK repeat
MBD: membrane-binding domain
SBD: spectrin-binding domain
ASD: autism spectrum disorder.

## Acknowledgements

Dr. Brenda R. Temple is gratefully acknowledged for initial modeling studies and many helpful discussions and advice. We thank Dr. Bryce W. Duncan for assistance and advice on mutagenesis and spine pruning assays. Luca Montore carried out initial AnkB/Spectrin binding assays.

---

Figure S1: Interaction Analysis of AnkB and NrCAM by PDBSum

PDBSum analysis of ANK repeats 1-24 of the AnkB MBD with the NrCAM cytoplasmic domain identified 8 salt bridges (red lines), 18 hydrogen bonds (blue lines), and 415 non-bonded contacts (orange dashed lines) between the indicated residues. Residue colors: positively charged (H,K,R-blue), negatively charged (D,E-red), neutral (S,T,N,Q-green), aliphatic (A,V,L,I,M-gray), aromatic (F,Y,W-purple), (P,G-orange). Arrows indicate the AnkB residues that were mutagenized and tested for NrCAM binding. The position of the 220 kDa molecular weight marker from PageRuler Plus is indicated and coincides with the AnkB220-HA band.

Figure S2: AnkB Associates with NrCAM in a Nexus of Residues in ANK Repeat R11

**Co-IPs**: Additional Western blots showing that NrCAM co-immunoprecipitates with WT AnkB-220 (HA-tagged) from transfected HEK293 cells, and to a much lesser extent with HA- AnkB-220 with point mutations H374W, H374A, R308E, and R297E.

AnkB was immunoprecipitated (IP) from cell lysates with anti-HA antibodies, and immunoblotted (IB) with anti-NrCAM antibodies followed by reprobing with anti-HA antibodies. NrCAM/AnkB ratios in the immunoprecipitates were obtained by densitometry relative to WT. Lanes from the same blots are shown without a line. Blots from different gels are shown with a wider separation. Nonimmune IgG did not pull down AnkB or NrCAM.

**Inputs:** Additional Western blots of WT and mutant HA-AnkB-220 in HEK293 lysates (5 µg) prior to immunoprecipitation as determined by immunoblotting (IB) with anti-HA antibodies, followed by stripping and reprobing with anti-NrCAM antibodies. Blots from different gels are separated. Levels of mutants relative to WT are shown below each lane. AnkB/Actin ratios of mutants relative to WT in inputs are shown (actin blots of lysates are shown in Fig. 3).

The position of the 220 kDa molecular weight marker from PageRuler Plus is indicated and coincides with the AnkB220-HA band.

Figure S3: Altered Interactions of ASD Missense Mutations AnkB A368G and A373V with NrCAM

**Co-IPs**: Additional Western blots showing that NrCAM co-immunoprecipitates with WT HA-AnkB-220 from HEK293 cells, and to lesser extent with HA-AnkB-220 with the mutation A368G. HA-AnkB-220 with mutation A373V shows increased binding to NrCAM. AnkB was immunoprecipitated (IP) from cell lysates with anti-HA antibodies, and immunoblotted (IB) with anti-NrCAM antibodies followed by reprobing with anti-HA antibodies. NrCAM/AnkB ratios in the immunoprecipitates were obtained by densitometry relative to WT on each gel. Lanes on the same blots are shown with or without a line.

Blots from different gels are shown with a wider separation. Nonimmune IgG did not pull down AnkB or NrCAM.

**Inputs:** Additional Western blots of WT and mutant HA-tagged AnkB in HEK293 lysates (5 µg) prior to immunoprecipitation as determined by immunoblotting (IB) with anti-HA antibodies, followed by stripping and reprobing with anti-NrCAM antibodies, and anti-Actin antibodies. Blots from different gels are separated. Levels of mutants relative to WT on each gel are shown under the lanes. The AnkB/Actin ratio of mutants relative to WT on each gel is shown below.

Figure S4: Interaction Analysis of AnkB and β2-Spectrin by PDBSum

PDBSum analysis of AnkB residues 966-1125 in the SBD with β2-Spectrin residues 1563-2093 in spectrin repeats 14 and 15 identified 2 salt bridges (red lines), 10 hydrogen bonds (blue lines), and 145 non-bonded contacts (orange lines) between the indicated residues. Residue colors: positively charged (H,K,R-blue), negatively charged (D,E-red), neutral (S,T,N,Q-green), aliphatic (A,V,L,I,M-gray), aromatic (F,Y,W-purple), (P,G-orange), (C-yellow). Arrow indicates AnkB R977, which is mutated in ASD variant AnkB R977Q.

Figure S5: Altered Interaction of ASD Missense Mutation AnkB R977Q with β2-Spectrin

**Co-IPs:** Additional Western blots of AnkB-220 co-immunoprecipitated with GFP-tagged β2-Spectrin from HEK293 cell lysates using anti-GFP antibodies. ASD mutation AnkB R977Q showed decreased association with β2-Spectrin, but AnkB P1380R was not affected. NrCAM/β2-Spectrin ratios in immunoprecipitates relative to WT on the same gels were obtained by densitometry. Lanes on the same blots are shown without separation lines. Blots from different gels are shown with a wider separation

**Inputs**: Additional Western blots of WT and mutant AnkB-220 in HEK293 lysates (5 µg) were determined by immunoblotting (IB) with anti-HA antibodies, followed by stripping and reprobing with anti-EGFP antibodies, anti-Actin antibodies or anti-Vinculin antibodies as loading controls. Levels of mutants relative to WT are shown below each lane.

Lanes from the same blot are shown without a line, or with a line if not adjacent. AnkB/Actin or AnkB/Vinculin ratios are shown below.

The position of the 220 kDa molecular weight marker from PageRuler Plus is indicated and coincides with the AnkB220-HA band.

Figure S6: Sema3F-Induced Spine Retraction is Impaired in Neuronal Cultures from *Ank2 ^+/-^* Mice Expressing the ASD Mutation AnkB A368G.

Raw images of *Ank2 ^+/-^* cortical neuronal cultures transfected with pCAG-IRES-EGFP and WT AnkB-220 or mutant AnkB-220 A368G are shown. Cells were treated on DIV14 with 5 nm Fc (control) or

Sema3F-Fc for 30 min, and immunostained for EGFP. Representative confocal images of EGFP-labeled apical dendrites on pyramidal neurons are shown. Scale bar = 3 µm.

