## Supplementary figures and images for "Structural Interactions of Ankyrin B with NrCAM and β_2_ Spectrin"

### Supporting Fig. S1

#### $AnkB_{1-24}$ – NrCAM interactions

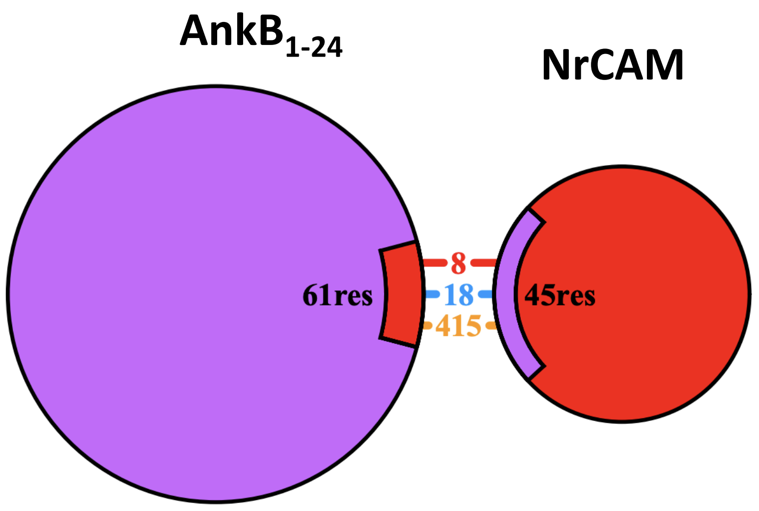


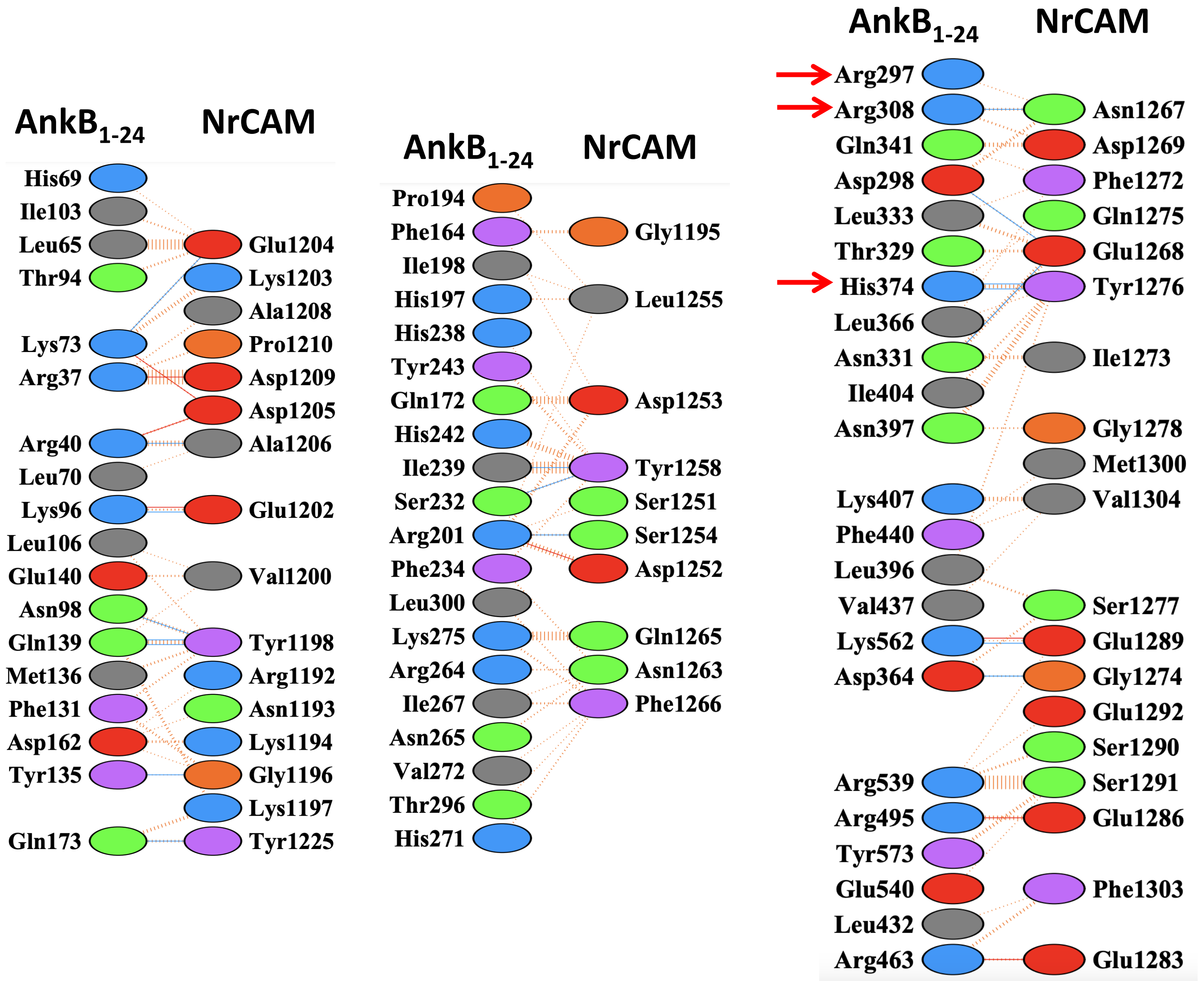

### Supporting Fig. S2

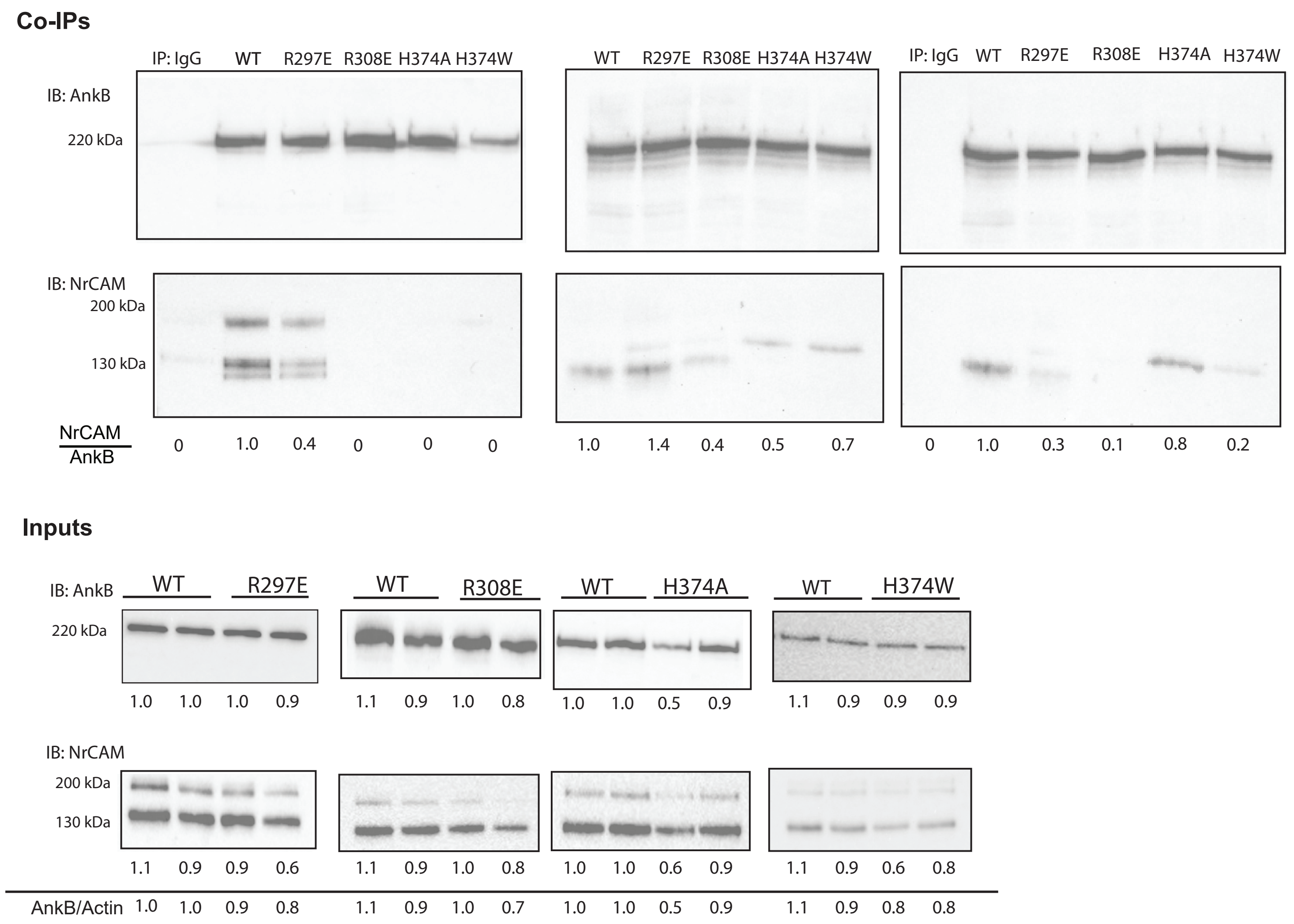

### Supporting Fig. S3

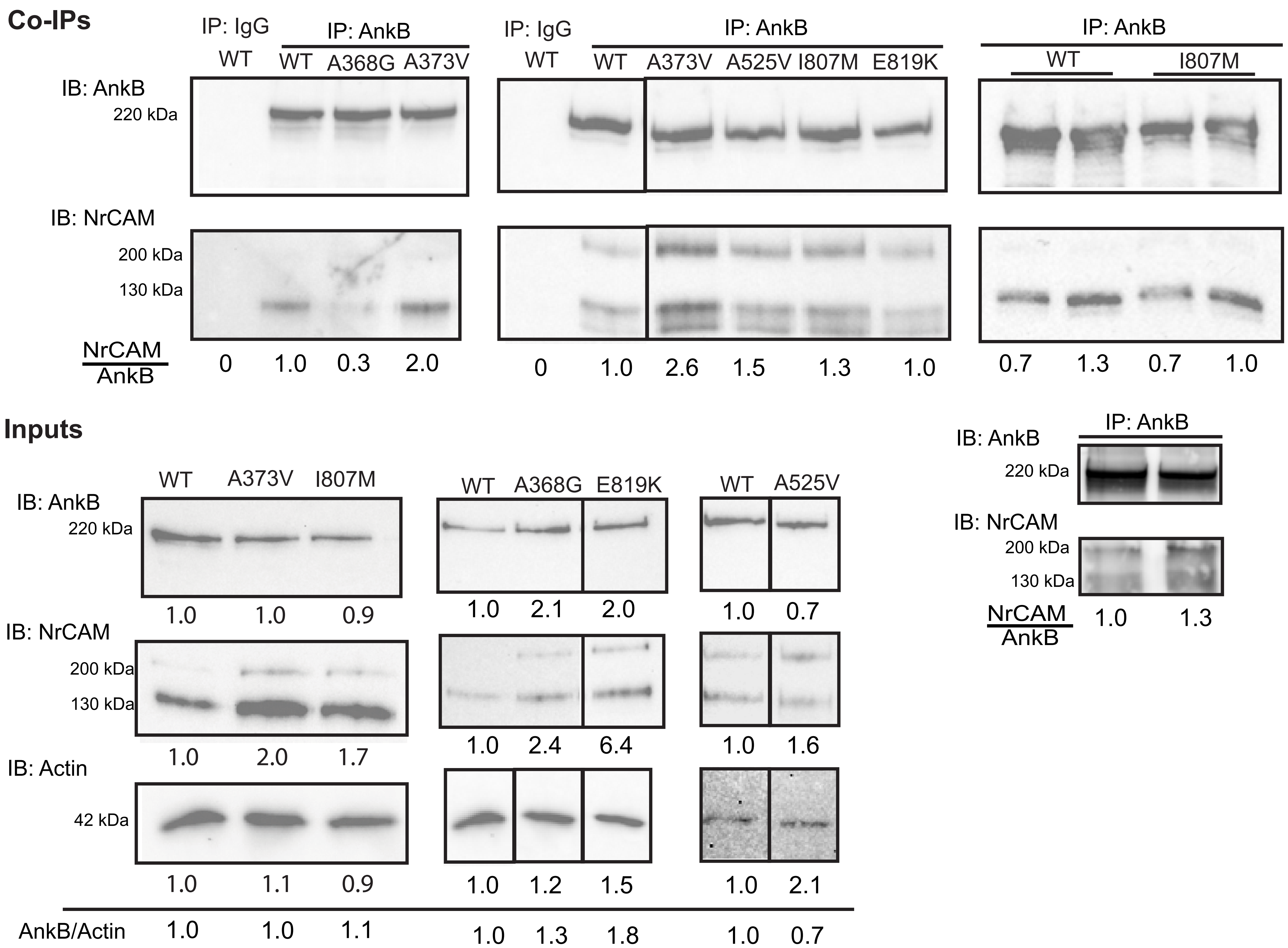

### Supporting Fig. S4

#### AnkB – β2-Spectrin interactions

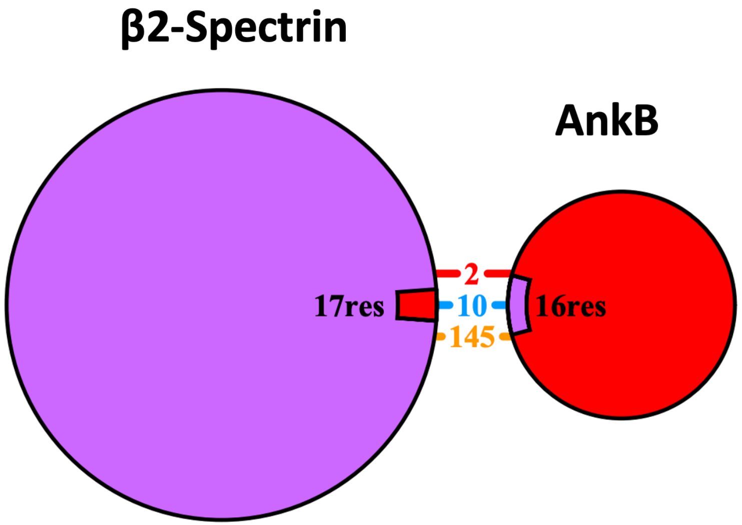


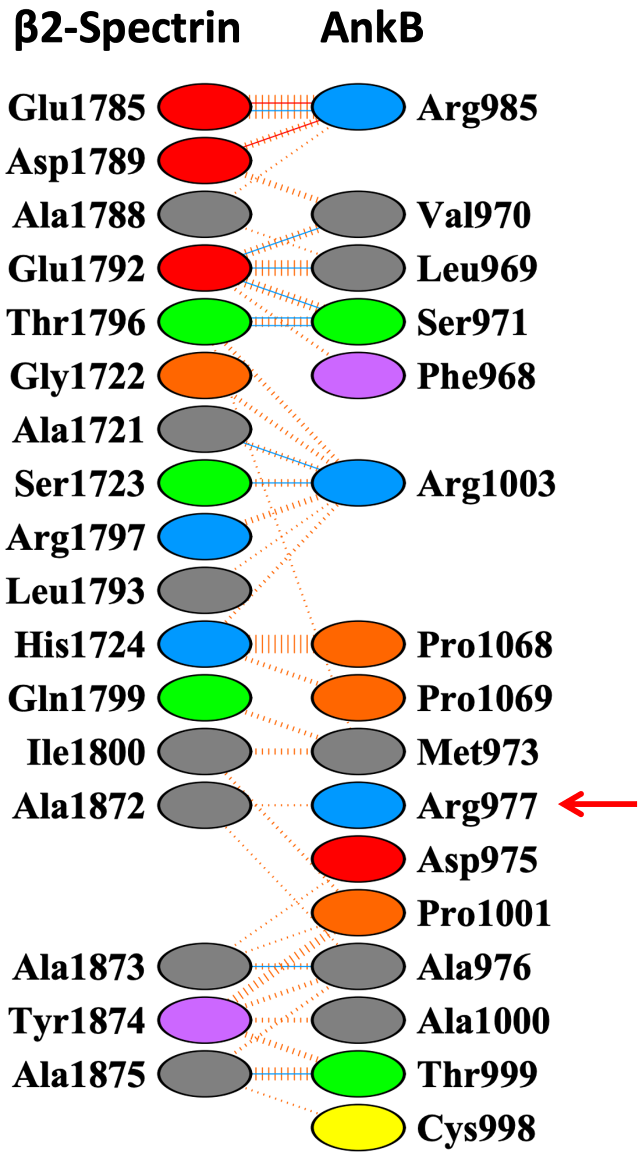

### Supporting Fig. S5

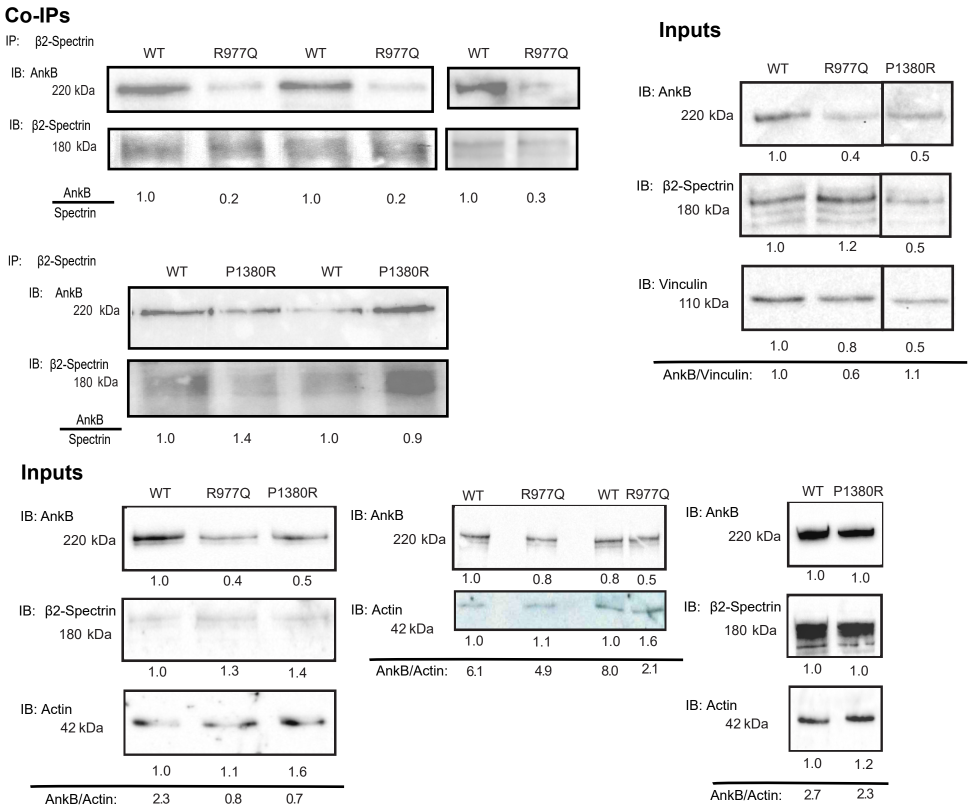

### Supporting Fig. S6

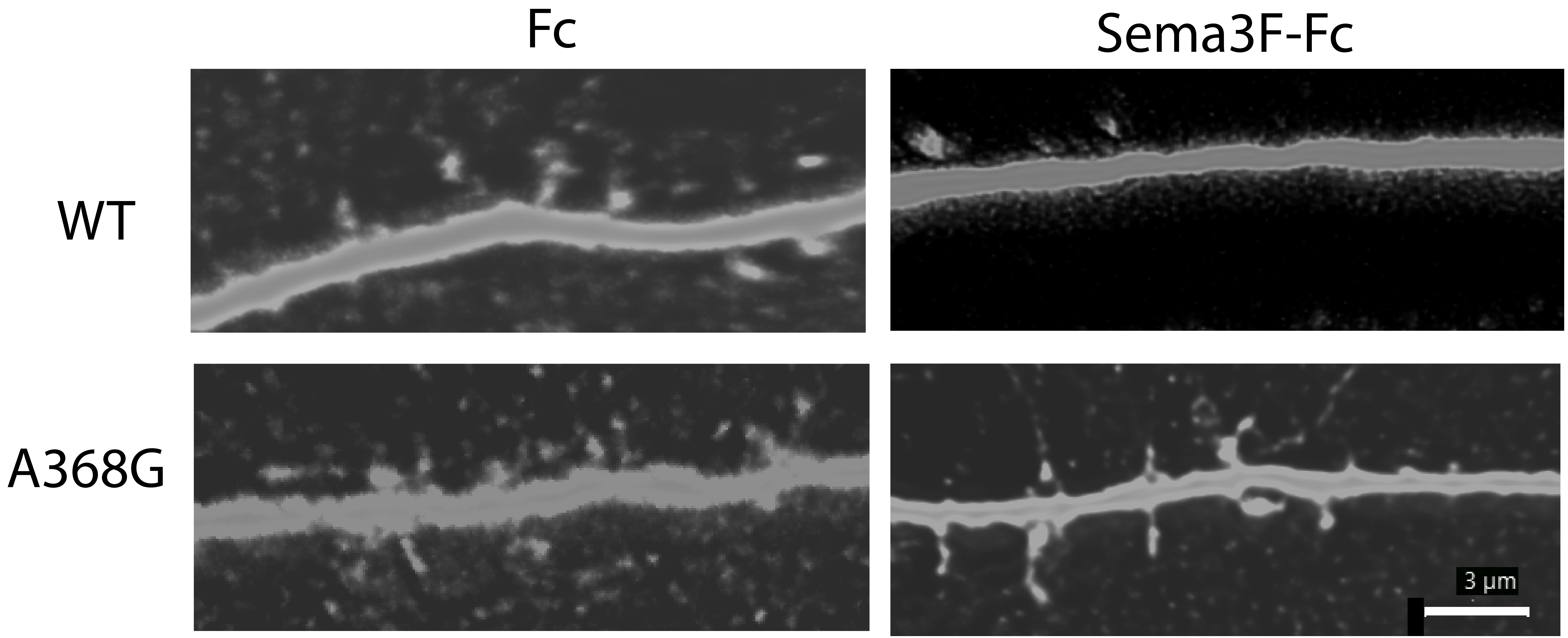
